# Automated detection of glaucoma with interpretable machine learning using clinical data and multi-modal retinal images

**DOI:** 10.1101/2020.02.26.967208

**Authors:** Parmita Mehta, Christine Petersen, Joanne C. Wen, Michael R. Banitt, Philip P. Chen, Karine D. Bojikian, Catherine Egan, Su-In Lee, Magdalena Balazinska, Aaron Y. Lee, Ariel Rokem, The UK Biobank Eye and Vision Consortium

**Author notes:** Members of the UK Biobank Eye and Vision Consortium include: Tariq Aslam, PhD, Manchester University, Sarah A. Barman, PhD, Kingston University, Jenny H. Barrett, PhD, University of Leeds, Paul Bishop, PhD, Manchester University, Peter Blows, BSc, NIHR Biomedical Research Centre, Catey Bunce, DSc, King’s College London, Roxana O. Carare, PhD, University of Southampton, Usha Chakravarthy, FRCOphth, Queens University Belfast, Michelle Chan, FRCOphth, NIHR Biomedical Research Centre,Sharon Y.L. Chua, PhD, NIHR Biomedical Research Centre, David P. Crabb, PhD, UCL, Philippa M. Cumberland, MSc, UCL Great Ormond Street Institute of Child Health, Alexander Day, PhD, NIHR Biomedical Research Centre, Parul Desai, PhD, NIHR Biomedical Research Centre, Bal Dhillon, FRCOphth, University of Edinburgh, Andrew D. Dick, FRCOphth, University of Bristol, Cathy Egan, FRCOphth, NIHR Biomedical Research Centre, Sarah Ennis, PhD, University of Southampton, Paul Foster, PhD, NIHR Biomedical Research Centre, Marcus Fruttiger, PhD, NIHR Biomedical Research Centre, John E. J. Gallacher, PhD, University of Oxford, David F. Garway-Heath, FRCOphth, NIHR Biomedical Research Centre, Jane Gibson, PhD, University of Southampton, Dan Gore, FRCOphth, NIHR Biomedical Research Centre, Jeremy A. Guggenheim, PhD, Cardiff University, Chris J. Hammond, FRCOphth, King’s College London, Alison Hardcastle, PhD, NIHR Biomedical Research Centre, Simon P. Harding, MD, University of Liverpool, Ruth E. Hogg, PhD, Queens University Belfast, Pirro Hysi, PhD, King’s College London, Pearse A. Keane, MD, NIHR Biomedical Research Centre, Sir Peng T. Khaw, PhD, NIHR Biomedical Research Centre, Anthony P. Khawaja, DPhil, NIHR Biomedical Research Centre, Gerassimos Lascaratos, PhD, NIHR Biomedical Research Centre, Andrew J. Lotery, MD, University of Southampton, Tom Macgillivray, PhD, University of Edinburgh, Sarah Mackie, PhD, University of Leeds, Keith Martin, FRCOphth, University of Cambridge, Michelle McGaughey, Queen’s University Belfast, Bernadette McGuinness, PhD, Queen’s University Belfast, Gareth J. McKay, PhD, Queen’s University Belfast, Martin McKibbin, FRCOphth, Leeds Teaching Hospitals NHS Trust, Danny Mitry, PhD, NIHR Biomedical Research Centre, Tony Moore, FRCOphth, NIHR Biomedical Research Centre, James E. Morgan, DPhil, Cardiff University, Zaynah A. Muthy, BSc, NIHR Biomedical Research Centre, Eoin O’Sullivan, MD, King’s College Hospital NHS Foundation Trust, Chris G. Owen, PhD, University of London, Praveen Patel, FRCOphth, NIHR Biomedical Research Centre, Euan Paterson, BSc, Queens University Belfast, Tunde Peto, PhD, Queen’s University Belfast, Axel Petzold, PhD, UCL, Jugnoo S. Rahi, PhD, UCL Great Ormond Street Institute of Child Health, Alicja R. Rudnikca, PhD, University of London, Jay Self, PhD, University of Southampton, Sobha Sivaprasad, FRCOphth, NIHR Biomedical Research Centre, David Steel, FRCOphth, Newcastle University, Irene Stratton, MSc, Gloucestershire Hospitals, NHS Foundation Trust, Nicholas Strouthidis, PhD, NIHR Biomedical Research Centre, Cathie Sudlow, DPhil, University of Edinburgh, Dhanes Thomas, FRCOphth, NIHR Biomedical Research Centre, Emanuele Trucco, PhD, University of Dundee, Adnan Tufail, FRCOphth, NIHR Biomedical Research Centre, Veronique Vitart, PhD, University of Edinburgh, Stephen A. Vernon, DM, Nottingham University Hospitals, NHS Trust, Ananth C. Viswanathan, FRCOphth, NIHR Biomedical Research Centre, Cathy Williams, PhD, University of Bristol, Katie Williams, PhD, King’s College London, Jayne V. Woodside, MRCOphth, PhD, Queen’s University Belfast, Max M. Yates, PhD, University of East Anglia, Jennifer Yip, PhD, University of Cambridge, and Yalin Zheng, PhD, University of Liverpool. These authors contributed equally (Aaron Y. Lee, Ariel Rokem).

## Abstract

Glaucoma, the leading cause of irreversible blindness worldwide, is a disease that damages the optic nerve. Current machine learning (ML) approaches for glaucoma detection rely on features such as retinal thickness maps; however, the high rate of segmentation errors when creating these maps increase the likelihood of faulty diagnoses. This paper proposes a new, comprehensive, and more accurate ML-based approach for population-level glaucoma screening. Our contributions include: (1) a multi-modal model built upon a large data set that includes demographic, systemic and ocular data as well as raw image data taken from color fundus photos (CFPs) and macular Optical Coherence Tomography (OCT) scans, (2) model interpretation to identify and explain data features that lead to accurate model performance, and (3) model validation via comparison of model output with clinician interpretation of CFPs. We also validated the model on a cohort that was not diagnosed with glaucoma at the time of imaging but eventually received a glaucoma diagnosis. Results show that our model is highly accurate (AUC 0.97) and interpretable. It validated biological features known to be related to the disease, such as age, intraocular pressure and optic disc morphology. Our model also points to previously unknown or disputed features, such as pulmonary capacity and retinal outer layers.

## Introduction

Glaucoma is the leading cause of irreversible blindness worldwide, affecting approximately 76 million people in 2020 and predicted to affect nearly 111.8 million by 2040 [76]. The risk of glaucoma increases with age [51]. As the older population grows due to longer life expectancy, glaucoma is expected to become a significant public health concern [76]. Glaucoma is characterized by the progressive loss of retinal ganglion cells, which manifests as thinning of the retinal nerve fiber layer (RNFL) and characteristic changes in the optic nerve head appearance [10]. In later stages of the disease, visual field defects develop and if uncontrolled, the disease can ultimately result in complete blindness.

Early detection and treatment of glaucoma is essential for minimizing risk of progressive vision loss and yet a number of challenges exist that prevent timely and accurate diagnosis. First, considerable expertise is required to perform the appropriate clinical exam and to interpret a number of specialized tests, such as visual field testing and retina and optic nerve imaging. The demand for this expertise is outpacing the supply of experts available to interpret tests and make diagnoses [20]. Second, glaucoma is often asymptomatic until the advanced stages of the disease. In the United States, approximately 50% of the estimated 3 million people with glaucoma are undiagnosed and in other parts of the world, estimates are as high as 90% [23, 68, 21, 17, 73]. New diagnostic tools that improve the diagnostic efficiency of the existing clinical workforce are therefore vital for enabling earlier detection of the disease to facilitate early intervention [28].

Although glaucoma is asymptomatic in its early stages, structural changes in the macula and RNFL precede the onset of clinically detectable vision loss [30]. Many studies have therefore attempted to automatically diagnose glaucoma using retinal imaging data. Most of these studies used either color fundus photos (CFPs) or features extracted from CFPs [7, 9, 67, 50, 12, 57, 63]. Other studies [49, 46] used features extracted from retinal B-scans obtained via Optical Coherence Tomography (OCT), a three-dimensional volumetric medical imaging technique used to image the retina. Macular OCT images are used to extract features such as thickness of the RNFL, ganglion cell-inner plexiform layer (GCIPL), or full macular thickness. Models evaluating changes in thickness of various retinal layers are promising since such changes, a direct result of tissue loss, are highly accurate disease predictors. However, thickness maps are derived automatically and, despite advances in OCT hardware and software, errors in segmenting retinal OCT images remain relatively common, with error estimates between 19.9% and 46.3% [44, 45, 5]. A study comparing a model built on raw macular OCT images with one built on thickness maps demonstrated that the former was significantly more accurate than the latter in detecting glaucoma [43].

In this work, we built a new, multi-modal, feature-agnostic model that includes clinical data, CFPs and macular OCT B-scans. Data for our model came from the UK Biobank, a multi-year, large-scale effort to gather medical information and data, with the goal of characterizing the environmental and genetic factors that influence health and disease [78]. About 65,000 UK Biobank participants underwent ophthalmological imaging procedures, which provided both macular OCT and CFP data that we matched with clinical diagnoses and with many other demographic, systemic and ocular variables. Specifically cardiovascular and pulmonary variables were chosen as markers of overall health. We used raw macular OCT and CFP data and did not rely on features extracted from these images. The use of machine learning, and particularly deep learning (DL), methods to analyze biomedical data has come under increased scrutiny because these methods can be difficult to interpret and interrogate [81, 6]; therefore, we applied machine learning interpretability methods to demystify and explain specific data features that led to accurate model performance [48]. Finally, we validated our model by comparing it to expert clinicians’ interpretation of CFPs to provide an additional benchmark for the performance of our machine learning model relative to current clinical practice.

## Methods

### Data access

Data was obtained through the UK Biobank health research program. Deidentified color fundus photos, OCT scans, and health data were downloaded from the UK Biobank repository and our study, which did not involve human subjects research, was exempt from IRB approval.

### Data set and Cohort selection

The UK Biobank is an ongoing prospective study of human health, for which data has been collected from over half a million individuals [71]. Participants throughout the UK were recruited between 2006 and 2010 and were aged 40-69 years at the time of recruitment. The data set contains information from questionnaires, multi-modal imaging measurements, and a wide range of genotypic and phenotypic assessments. Data collection is ongoing, allowing for longitudinal assessments. We analyzed a subset of the UK Biobank participants based on a snapshot of the repository that was created in the fall of 2017. This subset consisted of data from 96,020 subjects, 65,000 of which had retinal imaging data. This data set consisted of between one to three visits for each of the subjects. Color Fundus Photographs (CFP) data was available for only the first visit for these subjects. Retinal OCT data was available for first and second visits. The participants were given questionnaires to report various eye conditions, to which they could report healthy or chose one or more the following eye conditions: glaucoma, cataract, macular degeneration, diabetic retinopathy and injury for each eye. We used the answers provided as the labels for each eye. We did not examine the images to determine or alter the labels thus associated with the retinal image and clinical data.

#### Cohort selection

We selected a cohort from this data for the following three classes: A) subjects who in their first study visit report that they have been diagnosed with glaucoma and consistently report a glaucoma diagnosis in follow-up visits (glaucoma); B) subjects who in their first study visit report that they had no ocular conditions and consistently reported no ocular condition in follow-up visits (healthy); C) subjects who in their first visit report no ocular conditions, but in a subsequent visit report having received a diagnosis of glaucoma, labeled as the “progress to glaucoma” group (PTG). Ocular measurements were only available for the first two visits. The ocular data includes retinal imaging (both CFPs and macular OCTs) as well as IOP, Corneal hysteresis and Corneal resistance factor. However, a subset of the PTG group (n=21 retinas) received glaucoma diagnosis between the first and second visit and we used this subset to conduct statistical analysis of IOP. Systemic and pulmonary variables were available for the entire PTG group both pre- and post-diagnosis, and we were able to analyze the impact of diagnosis on these variables for the entire PTG group.

#### Exclusion criteria

We excluded all subjects who preferred not to answer questions about their ocular conditions, or did not know how to answer these questions. For *glaucoma* subjects, we excluded any subjects who listed any ocular conditions in addition to glaucoma, such as age-related macular degeneration, diabetic retinopathy, cataract, or injury. For the *healthy* subjects, we excluded any subjects whose visual acuity was recorded as worse than 20/30 vision in either eye. We also excluded any *healthy* subjects with any secondary health conditions (determined by primary diagnosis codes record in their hospital inpatient records). Finally, we excluded any retinal OCT scans from all three classes that could not be aligned using motion translation (x and/or y shift). Supplementary Figure 3. shows a sample of excluded retinal OCT images. The final number of for the three groups was glaucoma ((s)ubjects=863,(e)yes=1193), healthy (s=771, e=1283), and PTG (s=55,e=98). Supplementary Figure 4 shows the age and gender distribution of subjects in each of these groups. CFP images were available for only 56 of the 98 retinas in the PTG group (retinal OCT images were available for all PTG subjects).

#### Test set

At the outset, we randomly selected 100 eyes, 50 healthy and 50 with glaucoma. These were set aside as the test set on which we evaluated each of the models. We set an additional 170 eyes as a validation set for parameter tuning and model selection. The data was separated by subject such that both eyes of any subject belonged to either test, train or validation set. The test set was also rated by five glaucoma experts. Glaucoma experts used the CFPs for providing their scores. Glaucoma experts marked 13 CFPs from the test set as being such poor a quality to preclude any assessment. All comparisons of clinician and model performance excludes these 13 retinas. Supplementary Figure 5. shows a sample of excluded CFPs.

### Evaluating expert performance

Five glaucoma-fellowship trained ophthalmologists were recruited for the study to evaluate CFP images from test set to provide an expert diagnoses. The glaucoma experts identified the eye in each CFP as either healthy or glaucoma, and rated the confidence in the diagnosis from 1 to 5. A higher number indicated higher confidence in their diagnosis. This resulted in a 10-point scale for the diagnosis. We used this 10 point scale to create ROC curves for each expert.

### Machine learning models and training protocols

We built separate DL models for each imaging modality (retinal OCT and CFP). *Retinal OCT model* : the DL model built on the retinal OCT data took a single retinal OCT image as input and output a probability that the input image was from a subject with glaucoma. This model required individual B-scans. Each retinal OCT consisted of 128 B-scan images. This model was not provided any other additional information. This DL model was based on the densenet architecture [31], with four blocks with 6, 12, 48 and 32 layers each. We initialized model weights for this model with MSRA initialization [27]. Each retinal OCT B-scan is a gray scale 512 x 650 image. We flipped each right eye images left to right, we did this so that the optic nerve was on the same side for each scan. Additionally, we cropped each scan to an aspect ratio of 1:1 and down sampled to 224×224 pixels. We used a per pixel cross-entropy as the loss function with 0.1 label smoothing regularization [47]. We used Tensorflow [1] with Adam optimizer [36] and an initial learning rate of 1-e3 and epsilon of 0.1. We trained for 60 epochs(batch size 80) on one graphical processing unit (GPU)s. The hyper parameters for the training protocol were chosen by tuning on the validation data set. To improve the generalization ability of our model, we augmented the data by applying affine, elastic and intensity transformation over the input images.

#### CFP model

the DL model on the CFP took a single CPF image as input and outputs a probability that the input image was from a subject with glaucoma. This model was built with transfer learning [58, 2]. We chose transfer learning as (a) we had 128X fewer CFP images, and (b) CFP are color images and transfer learning has been shown to be effective for detecting other pathology in fundus images [40]. We used the InceptionResnetV4 [75] model, pre-trained on ImageNet data [32]. We used the Adam optimizer with an initial learning rate of 1-e5. We trained the model for 20 epochs, with a batch size of 400. During training, we kept the weights in 2*/*3 of the network (750 layers) frozen. We pre-processed each fundus image by flipping left CFP image so that optic nerve was on the same side of each image. We also subtracted local average color to reduce differences in lighting, and cropped the images to contain the area around the optical nerve (Supplementary Figure 5). We augmented the CFP by applying affine, elastic and intensity transformations similar to the retinal OCT images.

#### Baseline models

modern gradient boosted decision trees often provide state-of-the-art performance on tabular style data sets where features are individually meaningful, as consistently demonstrated by open data science competitions [22]. We used gradient-boosted decision trees, implemented in XGBoost [11], to build all three baseline models (BM1, BM2, BM3) based on demographic features: age, gender, ethnicity; systemic features: Body Mass Index (BMI), Forced Vital Capacity (FVC), Peak Expiratory Flow (PEF), heart rate, diastolic and systolic blood pressure, presence of diabetes, recent caffeine and nicotine intake; and ocular features: Intraocular pressure (IOP), corneal hysteresis, and corneal resistance factor. The systemic features were chosen as markers of overall health. We used the following hyper parameters for training: learning rate of 0.001, early stopping; 𝓁_2_ regularization of 1.0, no 𝓁_1_ regularization, no column sampling during training, and bagging sub-sampling of 70%. Hyper parameters were chosen by tuning on the validation data set.

#### Final model

we combined clinical data with results from image-based models to build the final model. To combine data from image models we used the probability of glaucoma as estimated by the respective image model as the feature value for each image. We combined these (128 OCT slices and one fundus) to a 129 element vector as the results of the image-based models. This vector was then combined with all of the features from BM3 for the final feature set. We used gradient-boosted decision trees to build this final model. The hyper parameters were chosen by tuning on the validation set and were as follows: learning rate 0.001, early stopping, bagging sub-sampling of 70%, 𝓁_2_ regularization of 1.0, no 𝓁_1_ regularization and no column sampling during training.

#### Interpretability Methods

for pixel-level importance in the image based DL models we used integrated gradients [72] and SmoothGrad [70] to determine salient pixels for the input images. For the tree-based models built using XGBoost, we used Tree explainer [42] to calculate the SHAP values. The SHAP values were used to determine feature importance and feature interaction.

#### Statistical analysis

We used bootstrapping [19] to determine confidence intervals for AUC and accuracy displayed in Figures 2 and 5. We performed analysis of variance(ANOVA) test to analyze the differences in pulmonary function features (FVC and PEF) among the three groups: healthy, glaucoma and PTG. We used the Dunn Test[18] with Bonferroni correction for pairwise comparison to determine differences between the three groups.

**Figure 1:**
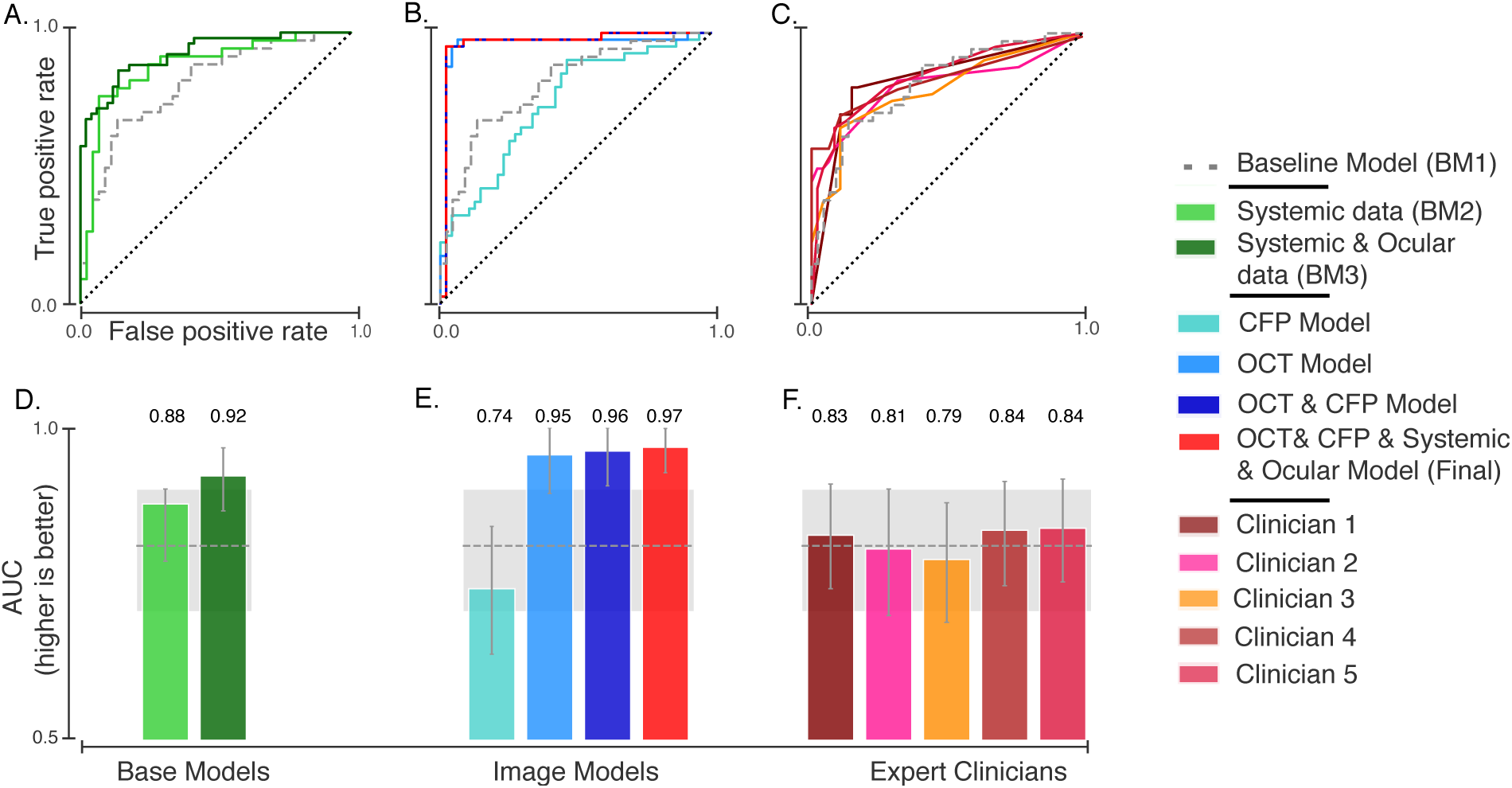
Results of glaucoma detection models. Receiver operating characteristic (ROC) curves are shown for (A) baseline models built with systemic and ocular data, (B) retinal imaging and final models, and (C) glaucoma expert ratings based on interpretation of CFPs. The corresponding area under the ROC curves (AUC) with (+/- 95% Confidence Interval) for models (D, E) and for clinician scores (F). The gray dashed line and shaded area denote the AUC and 95% CI for a base model (BM1) built on demographics (age, gender and ethnicity).

**Figure 2:**
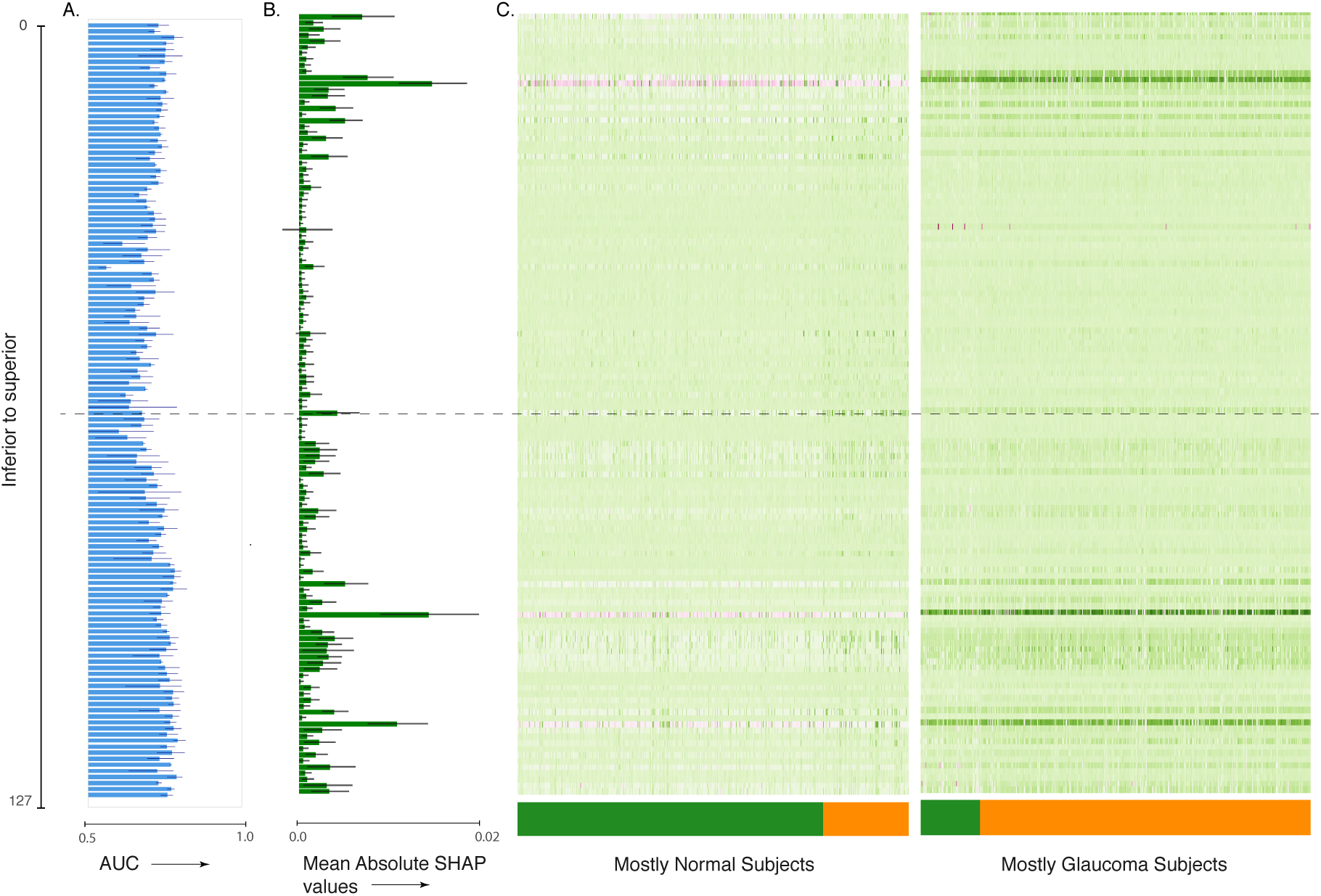
Interpretation of the ensemble model built on macular OCT images. **(A)** AUC for single image per retina models, **(B)** mean absolute SHAP values per retinal image for predicting glaucoma occurrence per retina, and **(C)** heat map of SHAP value per retinal image for predicting glaucoma occurrence per retina. The images are ordered from top to bottom and from superior to inferior retina. The dashed line indicates the central retinal image from the OCT volume.

**Figure 3:**
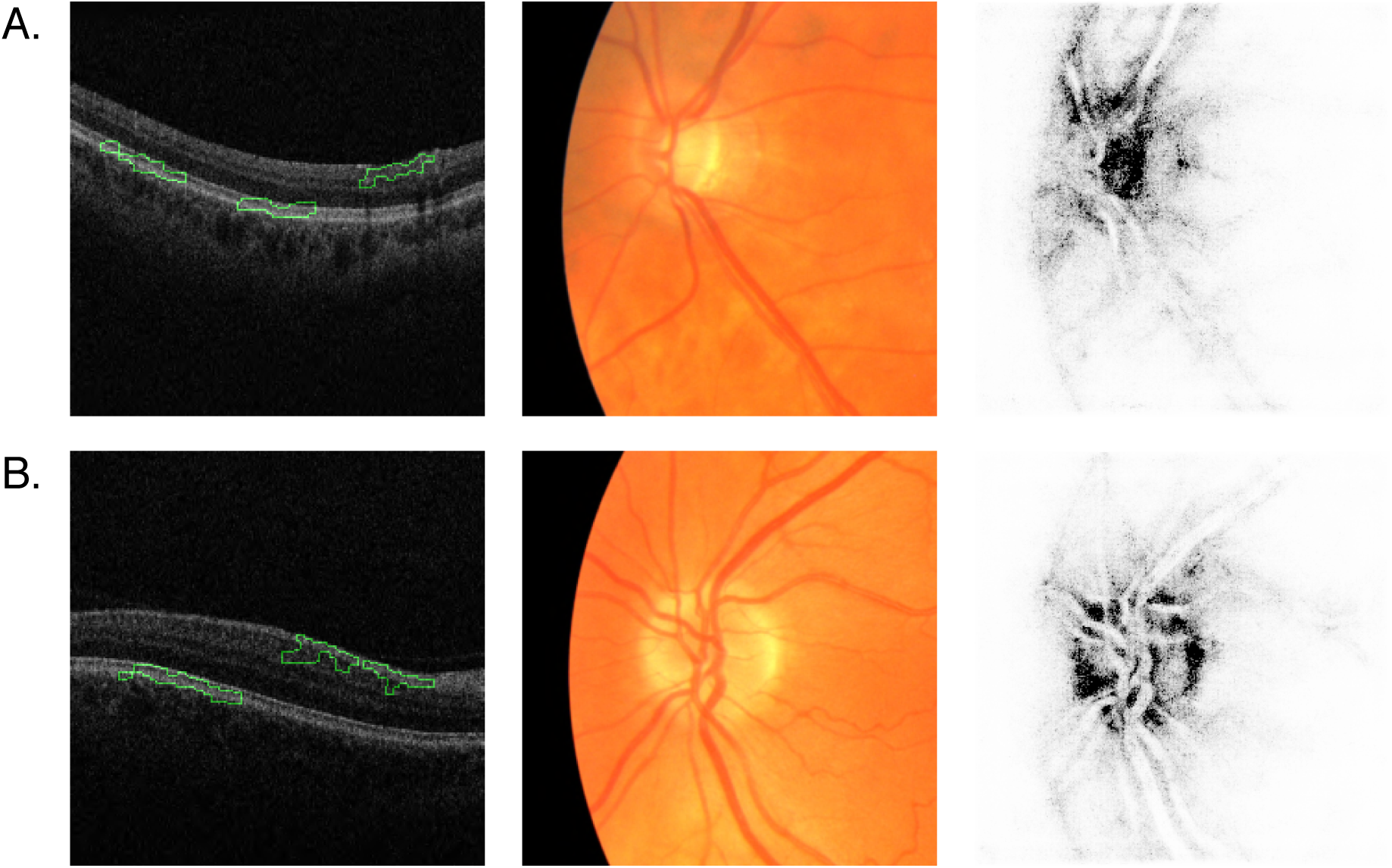
Saliency maps for macular OCT and CFP. Columns left to right: macular OCT image overlaid with saliency map, cropped CFP input to the neural network, CFP saliency map. Each macular OCT image is laid out with its temporal side to the left. **(A)** Retina of a subject with glaucoma diagnosis. **(B)** Retina of healthy subject. The green outline on the OCT saliency map indicates the areas the model deems most important. The darker pixels on the CFP saliency map indicate the areas the model deems most important.

**Figure 4:**
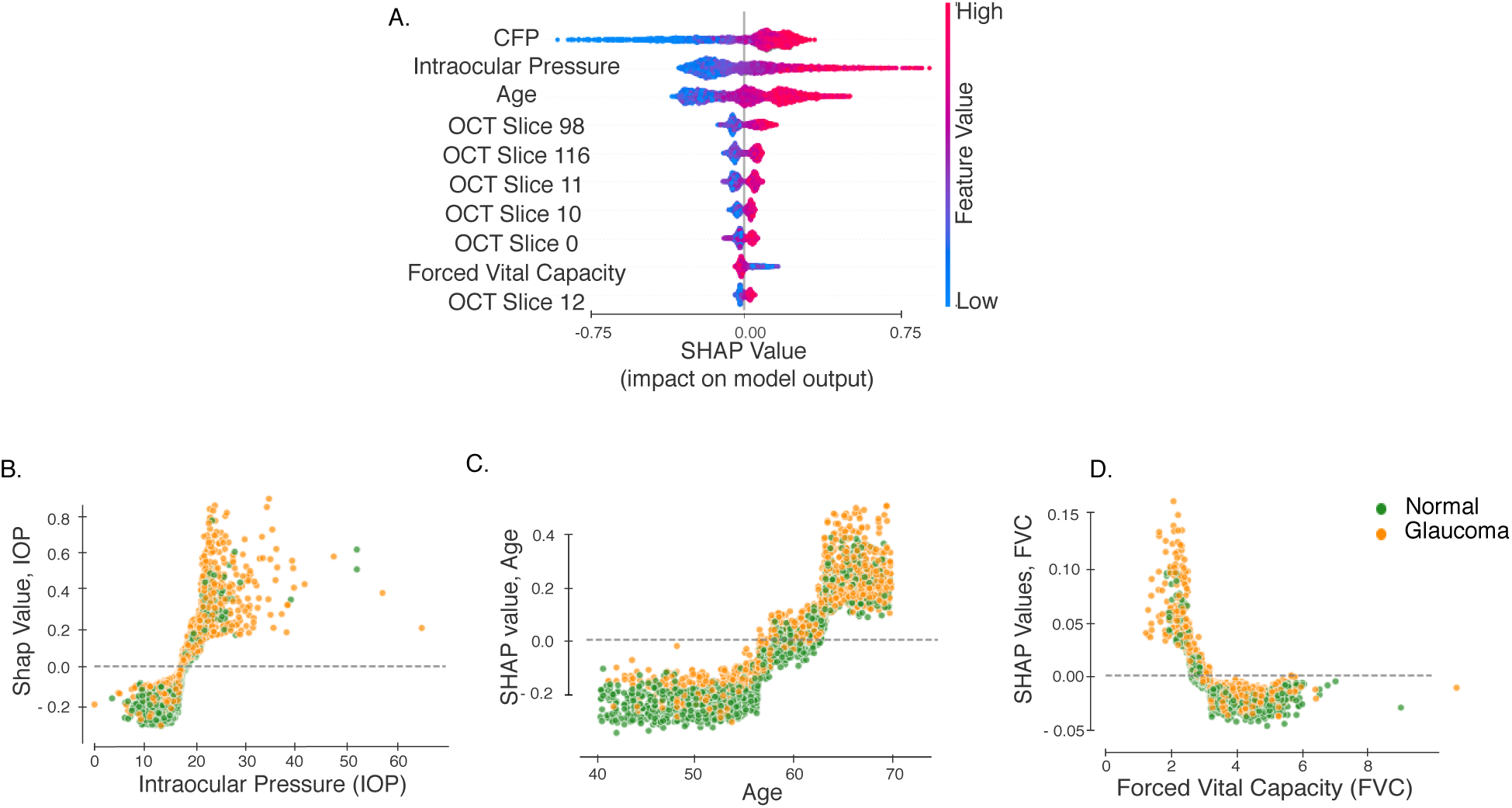
Interpretation of the final model built on image, demographic, systemic and ocular data. Interpretation for models built on medical and optometric features is based on SHAP values. **(A)** The 10 most important features from this model. SHAP values vs feature values for **(B)** Age, **(C)** IOP, **(D)** FVC and **(D)** BMI. Each point denotes a subject, and the color denotes whether the subject has been diagnosed with glaucoma.

**Figure 5:**
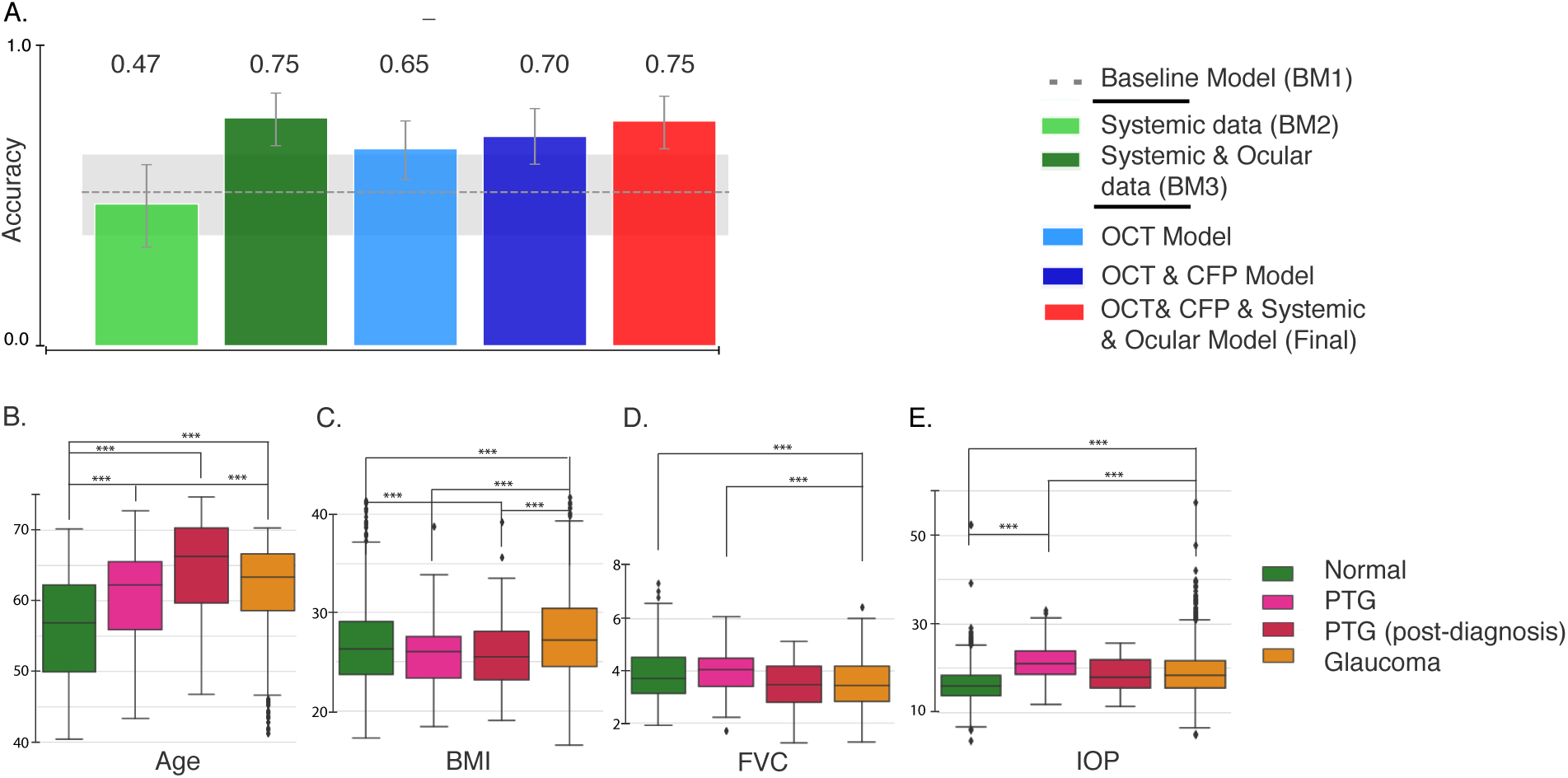
Evaluation of various models on the PTG cohort. (A) Accuracy of the models on the “progress-to-glaucoma” (PTG) cohort. The gray dashed line and shaded area denote the AUC and 95% CI for a base model built on demographics alone (age, gender and ethnicity; BM1). The bottom row shows the distribution of Age(B), BMI(C), FVC(D) and IOP(E) for healthy, PTG, PTG (after glaucoma diagnosis) and glaucoma. ***P < 0.0001.

#### Code availability

The code used to fit, evaluate and interpret the models is available at: https://github.com/uwdb/automatedglaucomadetection.

## Results

### We built multiple models using clinical data to establish a baseline

Glaucoma is related to many biological features, the most important of which is age [24]. Thus, we built our first baseline model (BM1) on basic demographic characteristics of the patient and control populations. BM1 included age, gender and ethnicity. Using these features, a boosted gradient tree based model predicted an occurrence of glaucoma well above chance (area under the ROC: 0.81, 95% CI 0.71-0.90). In addition, we created two other models. The systemic data model (BM2) added cardiovascular and pulmonary variables – including Body Mass Index (BMI), Forced Vital Capacity (FVC), Peak Expiratory Flow (PEF), heart rate, diastolic and systolic blood pressure, and the presence of diabetes – to the demographic variables from BM1. We also included transient factors, such as recent caffeine and nicotine intake, to account for any transient impact on blood pressure and heart rate. BM2 outperformed BM1 (0.87 AUC, 95% CI: 0.79-0.96). In the third model (BM3), we added ocular data to BM2, including IOP, corneal hysteresis, and corneal resistance factor. BM3 outperformed both of the other baseline models, with a test set AUC of 0.92, 95% CI: 0.87 - 0.96; see Figure 1A,D.

We used SHapley Additive exPlanations (SHAP) [41] to analyze the features that provide high predictive power in BM3. SHAP allocates optimal credit with local explanations using the classic Shapley values from game theory [66] and provides a quantitative estimate of the contribution of different features to the predictive power of a model. A higher absolute SHAP value indicates greater feature impact on the model prediction and greater feature importance. The five features with the highest mean absolute SHAP values for BM3 were age, IOP, BMI, FVC and PEF. Supplementary Figures 1 and 2 show the most important features in BM3, as evaluated through SHAP, and the interaction effect among the top features.

### We built a separate DL model on each retinal image modality

Glaucoma is characterized by structural changes in the optic disc and other parts of the retina. Visual examination of CFP and macular OCT images is therefore an important tool in current diagnostic practice [52]. Since our data set included both CFP and OCT images, we built separate DL models for each image modality (see Methods). The DL model built on CFP classified eyes diagnosed with glaucoma with modest accuracy (AUC: 0.74, 95% CI: 0.64-0.84; Figure 1B,E). The DL model built on macular OCT images was more accurate than all of the baseline models and the model trained on CFP images (AUC: 0.95, 95% CI: 0.90-1.0). When we combined information from both the DL models trained on CFP and OCT via an ensemble, the resulting model was marginally more accurate than the DL model build on macular OCT images alone (AUC: 0.963, 95% CI:0.91-1.0). This suggests that most CFP information is redundant with information present in macular OCT images.

### We used several methods to interpret the DL models

DL models are notoriously inscrutable. However, several methods for interrogating these models have recently emerged [74, 55, 56, 69, 72]. To assess the features that lead to high performance of the image-based models, we first assessed which scan of the macular OCT provided the most information. We fit individual models to each scan of the macular OCT. Recall that macular OCTs are volumetric images; in the UK Biobank data set, each macular OCT consists of 128 scans. We found that models using scans from the inferior and superior macula were more accurate than those using the central portion of the macula (Figure 2A). Second, we built an ensemble model that used the results of the DL models of the individual macular OCT scans to predict glaucoma occurrence per retina. This model used each of the 128 macular OCT scans to make a prediction about the retina. Figure 2B shows the feature importance attributed to each scan via SHAP; it shows that scans from the inferior retina were deemed more important by this model. Large patient and control populations are heterogeneous, and we do not generally expect that information will consistently come from one particular part of the retina. Nevertheless, when considering the SHAP values of each macular OCT scan, we found that the data set broke into two major clusters based on the SHAP values from different retinal parts (Figure 2C). One cluster mostly contained retinas from healthy subjects and used scans from the inferior part of the retina as negative predictors of glaucoma. The second cluster mostly contained glaucomatous retinas, and SHAP values of these same scans from inferior and superior macula were used as positive predictors of glaucoma. This also explains why models fit only to scans from the inferior or the superior macula were more accurate.

In addition to the scan-by-scan analysis, image-based models can be evaluated pixel-by-pixel to determine the importance of specific image features to the DL models’ decision making. Using integrated gradients [72], we generated saliency maps of the pixels responsible for DL model prediction. Figure 3 shows a macular OCT scan for an eye with glaucoma and a scan for a control eye along with the CFP images and CFP saliency maps for each eye. The CFP saliency maps typically highlight the optic nerve head in both normal and glaucomatous retinas. The saliency maps for OCT image typically highlight the nasal side of the RNFL and outer retinal layers.

### We built the final model by combining both modalities of retinal imaging, demographic, systemic and ocular features

This model was an ensemble, which combined information from raw macular OCT B-scans and CFP images as well as all demographic, systemic and ocular data used in BM3. This final model had an AUC of 0.967, (95% CI: 0.93 - 1.0). Figure 4 shows the ten features with the highest mean absolute SHAP value over all observations in the data set. The most important features for this final model, as determined by their SHAP values, include age, IOP and FVC, in addition to the CFP and macular OCT scans from both inferior and superior macula. BMI is less significant than FVC in this final model. Further, IOP had a higher importance than age. This is a reversal in importance of features when compared to models built without information from retinal imaging. Unsurprisingly, this confirms that the CFP and OCT scans contain information that supersedes in importance the information provided by BMI and age.

### We compared the performance of our model with ratings from glaucoma experts to provide a comparison to current clinical practice

To compare the performance of our final model to expert clinicians, five glaucoma experts evaluated CFPs of the test set. Initially, experts were also given access to OCT images for each subject. However, raw b-scans from macular OCTs are not an image modality that experts usually examine during regular clinical practice for glaucoma diagnosis. Since we did not have access to thickness maps, experts made the diagnoses using only the CFP data. (Figure 1C and D). The highest AUC for the expert rating was 0.84, and the lowest was 0.79. The average pairwise kappa for the five experts was 0.75, indicating a good level of agreement between experts about the diagnosis.

### We validated our model by evaluating it on patients that progress to glaucoma

The UK Biobank data set contained several subjects who lacked a glaucoma diagnosis on their first study visit but received a diagnosis before a subsequent visit. These “progress-to-glaucoma” (PTG) subjects provide a unique opportunity to evaluate our model, which was built on data from glaucomatous and healthy subjects. Detection of glaucoma in the PTG cohort was tested using all of our models (Figure 5A). Both BM1 (based on age, gender and ethnicity) and BM2 (added systemic variables) were indistinguishable from chance performance (BM1: 51 % correct: 95% CI [36% - 64%]; BM2: 47% correct: 95% CI[33%-60%]). BM3, which included ocular variables, achieved substantially higher accuracy at 75% correct (95% CI: [67%-83%]). The model trained on macular OCT images achieved slightly lower accuracy at 65% correct (95% CI [55% -74.5%]), and the model trained on combined CFP and OCT achieved an accuracy of 69% (95% CI [60.2% -78.6%]). The final model trained on OCT, CFP and all other available features achieved an accuracy of 75% correct (95% CI [65% - 83%]).

Since PTG subjects did not undergo glaucoma treatment, this evaluation provides additional insight into the biological features of the disease. For many of these features, including age and BMI, the PTG group lie between the normal and glaucoma groups (Figure 5B to E). We identified two interesting deviations from this pattern. First, for the pulmonary capacity variables (FVC and PEF), the PTG group was indistinguishable from the healthy subjects in our sample, and both healthy and PTG subjects significantly differed from patients with glaucoma. This difference is statistically significant even when controlling for age (see Supplementary Results). However, on a subsequent visit, after receiving a glaucoma diagnosis, the pulmonary capacity measurements of this group was indistinguishable from that of the glaucoma group. Second, the PTG group had a significantly higher IOP than the group diagnosed with glaucoma (Figure 5D; see Supplementary Results). The post-diagnosis IOP measurements of the PTG group shows similar trend with lower IOP values.

## Discussion

Automating glaucoma detection using imaging and clinical data may be an important and cost-effective strategy for providing population-level screening. In this study, we used machine learning to construct an interpretable machine learning model that combined clinical information with multi-modal retinal imaging to detect glaucoma. We created and compared several models based on clinical data to establish a baseline: BM1 used demographic data (age, gender, ethnicity), BM2 added systemic medical data (cardiovascular, pulmonary), and BM3 added ocular data (IOP, corneal hysteresis, corneal resistance factor). Our final model was an ensemble, which combined information from raw macular OCT B-scans and CFP images as well as all demographic, systemic and ocular data used in BM3. This final model had an AUC of 0.97.

In interpreting this final model, we found that CFP, age, IOP, macular OCT images from the inferior and superior macula, and FVC were the most important features (Figure. 4). The significance placed upon age and IOP by our final model reiterate previously known risk factors for glaucoma. The positive SHAP values for IOP in our model rapidly increased above an IOP of approximately 20, consistent with the fact that ocular hypertension is a key risk factor for the disease [33, 62, 25, 3, 65]. Age and IOP switched places in their relative importance in our final model, which includes retinal imaging, in addition to BM3 features. This suggests that retinal imaging includes information that supersedes or is redundant with information linked to age. This finding is consistent with previous research, which demonstrated the ability of CFP to predict cardiovascular risk factors including age [64]. We also observe two discontinuities in the age vs. SHAP values for age (Figure. 4B), at ages 57 and 65. At both of these ages the SHAP values for age increase at a higher rate than before. This could be both due to biological, as well as socio-economic factors (e.g., 65 is the age of retirement in the UK).

An important novel finding of our study was the correlation of pulmonary measures, especially decreased FVC, with glaucoma. There are several possible explanations for this finding. First, a recent study by Chua et al. found a correlation between glaucoma and atmospheric particulate matter [15]. Chua et al.’s study did not include pulmonary function tests such as FVC and was correlational in nature but other studies have linked exposure to particulate matter with decreased FVC [13, 80, 26], suggesting common causes for reduced FVC and for glaucoma. Second, it may be that the treatment of glaucoma with topical beta blocker therapy has an impact on reducing FVC [61]. This idea receives further support from the findings in the PTG group, who do not have a diagnosis and have presumably not received any treatment. These individuals have FVC that is higher than the glaucoma group and is indistinguishable from the healthy group before a diagnosis is made. After a diagnosis is made, their FVC also decreases to levels indistinguishable from that of the glaucoma group. Thus, lower FVC values could indicate a result of glaucoma treatment.

Examination of the pixel-by-pixel importance of both retinal image modalities provided additional insight into what our model focused on when predicting glaucoma (Figure 3). For the CFPs, the model focused on the optic disc, a known source of information in the clinical diagnosis of glaucoma [4]. For the macular OCT B-scans, the model relied on previously validated retinal areas, including the inferior and superior macula [60]. In addition, the algorithm points to the nasal macular RNFL. The effect of glaucoma on RNFL integrity is well-understood, and RNFL thickness maps are often used clinically to diagnose glaucoma. However, the automated algorithms that are used clinically have a high segmentation error rate, resulting in variable thickness estimates, which may in turn lead to errors in diagnosis [44]. By avoiding reliance on extracted features such as thickness maps, our approach enabled the discovery of other possible biological features of glaucoma. For example, consistent with recent results in the same data set [35], the model also identified other (non-RNFL) parts of the inner retina as important (e.g., see Figure 3B).

In addition to the RNFL and inner retina, the model relied on the outer layers of the retina for glaucoma diagnosis. The involvement of the retinal outer layers in glaucoma is controversial. In a typical analysis of OCT images that focuses on the thickness of different parts of the retinal layers, glaucoma effects are usually not found in outer layers [79, 16, 37], but an association between age, IOP and retinal pigment epithelium thickness is sometimes detected [38]. Some anatomical studies do not find any differences in the outer retinal layer between healthy and glaucomatous eyes [34]. Other studies, using psychophysical methods in human subjects with glaucoma [29, 54], using histological methods in human eyes [59, 53] or examining animal models of glaucoma [8, 53] have shown the involvement of the retinal outer layer in glaucoma. In addition, Choi et al. [14] used high-resolution imaging techniques (ultrahigh-resolution Fourier-domain optical coherence tomography, UHR-FD-OCT, and adaptive optics, AO) to image glaucomatous retinas. They found a loss of retinal cone receptor cells that correspond to visual field loss. This loss of cones could cause subtle changes in the appearance of this part of the retina, that are not reflected in changes in thickness but are still captured by the DL model (e.g., changes in texture). The ability of DL models to use visual cues that are not apparent to the human eye has been previously demonstrated in another study in which retinal angiograms were generated from OCT images [39]. This finding is also consistent with a recent study that used unsegmented OCT scans and reported the involvement of outer retinal layers in a DL model that detects glaucoma [43, 77].

Our final model detected the occurrence of glaucoma with an accuracy of 75% on a cohort that had not yet been clinically diagnosed at the time of their testing (“progress-to-glaucoma", PTG). This does not constitute early detection: even though the individuals were not clinically diagnosed, they may already have significant disease progression, since many patients are undiagnosed even in relatively late stages of the disease [73]. The median IOP value was higher for the PTG cohort than for the subjects diagnosed with glaucoma, possibly because treatments for glaucoma are designed to decrease IOP. The PTG group also tended to be younger than those diagnosed with glaucoma. Interestingly, FVC in the PTG group was higher than in the glaucoma group and was indistinguishable from healthy subjects. This finding helps explain why BM2, which relied heavily on PVC and PEF, performed relatively poorly on the PTG cohort, achieving an AUC of 47% (Figure 5A). It also provides possible evidence against a causal relationship between FVC and glaucoma, as mentioned above. Furthermore, in a post-diagnosis visit, pulmonary factors (FVC and PEF) in these individuals were lower and indistinguishable from that of the patients with glaucoma, further supporting a possible treatment effect. This area warrants further investigation.

Before our model can be considered for use in a real-world setting, several limitations should be considered and addressed. First, we included only subjects without other ocular disorders. In the general population, glaucoma may coexist with other ocular comorbidities, and it is unclear what effect this may have on the model’s ability to detect glaucoma accurately. However, selecting subjects with only a glaucoma diagnosis and no other ocular morbidities instills confidence that the model we built is glaucoma-specific, delineates the boundaries between these groups, and identifies the features specific to glaucoma. Second, features of the optic disc are clinically important in diagnosing glaucoma. The limited quantity and poor quality of the CFPs in the UK Biobank data set likely contributed to the low AUC of both the CFP DL model and the expert clinician grading. Finally, a major limitation of our study was the veracity of ground truth labels used to train the model. Labels used were based on self-report. While we eliminated any subject who had inconsistent answers or declined to answer, the generally high rate of undiagnosed glaucoma and the potential for recollection error means that some participants may have been incorrectly labeled.

Our study combined information from multiple sources – including two different retinal imaging modalities (CFP and OCT), demographic data, and systemic and ocular measurement – to build a model that detects glaucoma. This approach yielded not only very accurate detection, but it also enabled us to isolate and interpret critical variables that helped us draw clinical insights into the pathogenesis of the disease.

## Acknowledgments

AR was supported through a grant from the Betty & Gordon Moore Foundation and the Alfred P. Sloan Foundation to the University of Washington eScience Institute Data Science Environment. AYL was support by NIH/NEI K23EY029246 and an unrestricted grant from Research to Prevent Blindness and the Lowy Medical Research Institute. PM and MB were supported by NSF grants AITF 1535565, NSF grants IIS 1524535 and a gift from Intel. SL was supported by NIH R35 GM 128638.

## Author contributions

PM, MB, AR and AYL initiated the study. PM, SL, MB, AYL and AR designed the computational analysis. AYL provided computational resources. PM implemented the computational analysis and conducted the analysis. CE and the UK Biobank Eye and Vision Consortium collected and provided access to the UK Biobank data. CP, JCW, MRB, PPC and KDB examined and diagnosed retinal imaging data. PM, CP, AYL and AR wrote the manuscript. All authors provided manuscript feedback.

## 1 Supplementary Results

**Supplementary Figure 6:**
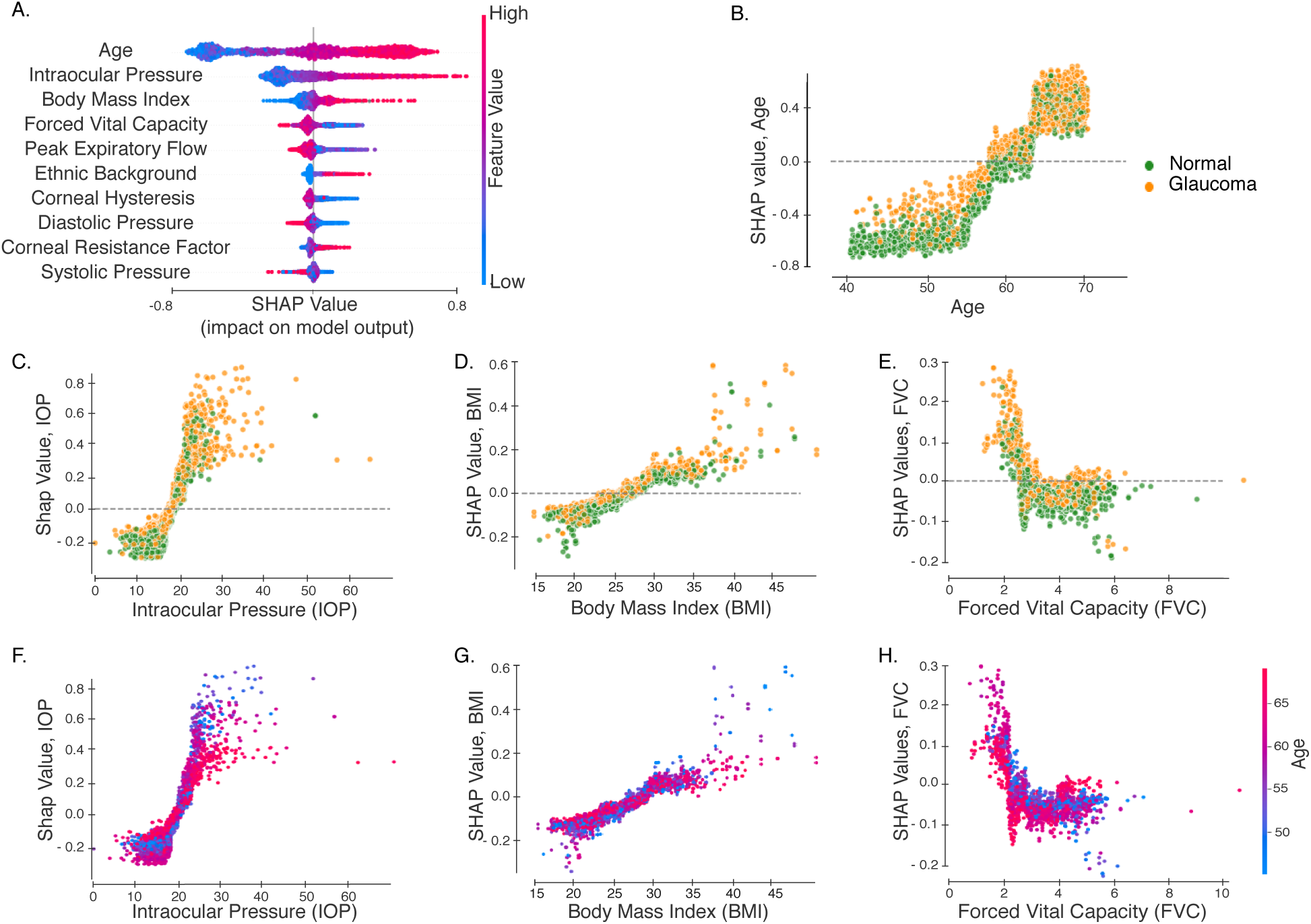
Interpreting model built on demographic, systemic and ocular data. A) The ten most important features from BM3 based on SHAP values. B-E) SHAP values vs feature values for age, IOP, BMI, and FVC respectively. Each point represents an individual subject and the color denotes whether or not they have been diagnosed with glaucoma. F-H) SHAP values vs feature values for features IOP, BMI, and FVC respectively, with each point colored based on age of subject.

**Supplementary Figure 7:**
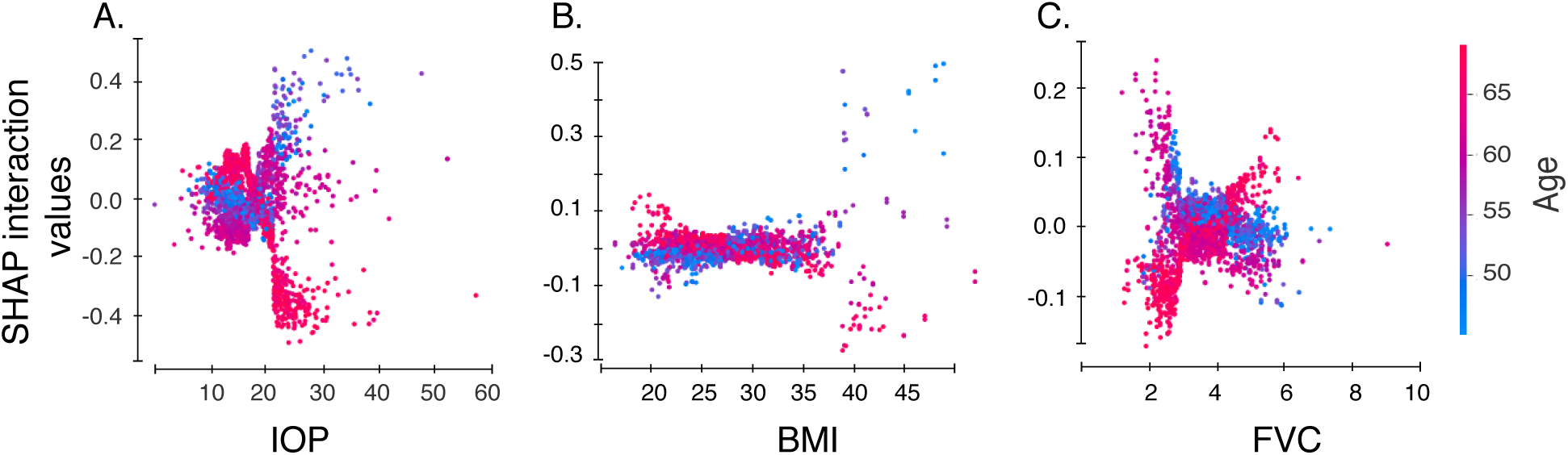
SHAP interaction values in BM3. Interactions between age and each of the other top variables in model BM3: **A** SHAP interaction values for age and IOP. **B** SHAP interaction values for age and BMI. **C** SHAP interaction values for.

### 1.1 Statistical analysis of group differences

Statistical analysis of differences between glaucoma, healthy, PTG and PTG post-diagnosis demonstrates that pulmonary capacity variables differ between the groups. This is true even when controlling for age: an ANCOVA of FVC values for the three groups, using age as a covariate found a statistically significant effect of group identity (Supplementary Table 1). To further confirm this, we followed this ANCOVA with a pairwise Dunn’s test (Supplementary Table 2 and Supplementary Table 3). We found that FVC in PTG significantly differed from FVC in patients with glaucoma (Supplementary Table 2). Likewise, FVC in normal vs. patients with glaucoma differed significantly (Supplementary Table 2), but we did not find a difference between normal and PTG (Supplementary Table 2). Values for PEF were similar: the ANCOVA using age as a covariate showed a statistically significant effect of group (Supplementary Table 1); Dunn’s tests showed similar differences: glaucoma differ significantly from both PTG and healthy, but healthy and PTG do not significantly differ (Supplementary Table 3). In addition, a pairwise Dunn’s test confirmed that the IOP difference between glaucoma and PTG groups was statistically significant (Z=-10.74, p<0.05). Post-diagnosis, the difference between PTG and glaucoma in FVC and PEF was no longer statistically significant. This was also true for IOP, but this is based on only a limited sample of PTG (n=21 retinas), for which there is a measurement of IOP post-diagnosis (see Methods).

**Supplementary Table 1:** Analysis of covariance (ANCOVA) of pulmonary capacity variables (forced vital capacity (FVC), peak expiratory capacity(PEF)) for group identity amongst the three groups (Healthy, Glaucoma and PTG), controlling for age.

| Dependent Variable | dF | F Value | p value |
| --- | --- | --- | --- |
| FVC | 1,2403 | 64.51 | $1.49e-15$ |
| PEF | 1,2403 | 50.8 | $1.35e-12$ |

**Supplementary Table 2:**
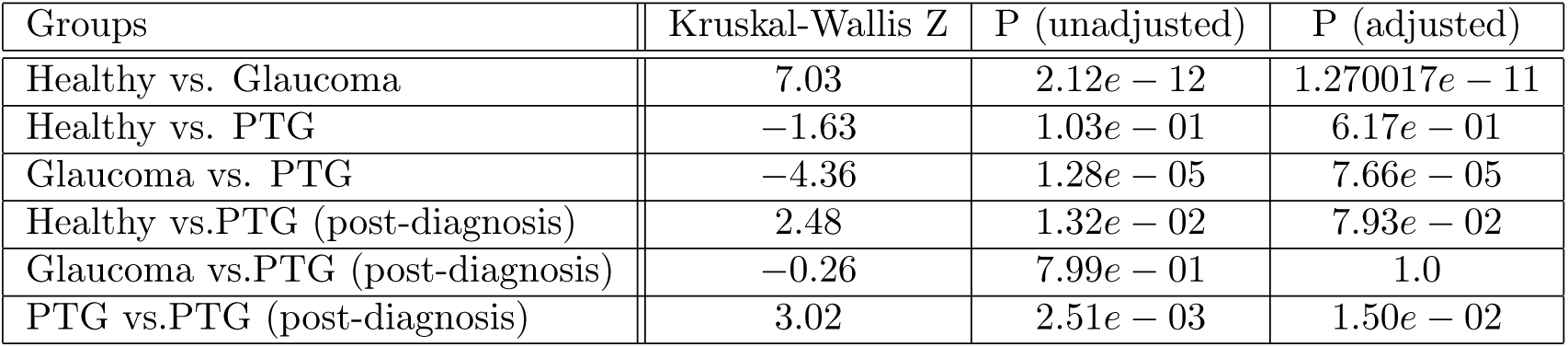
Dunn’s test comparing **FVC** for Glaucoma, healthy, PTG and PTG (post-diagnosis) groups, with Bonferroni correction for p-values.

| Groups | Kruskal-Wallis Z | P (unadjusted) | P (adjusted) |
| --- | --- | --- | --- |
| Healthy vs. Glaucoma | 7.03 | $2.12e-12$ | $1.270017e-11$ |
| Healthy vs. PTG | -1.63 | $1.03e-01$ | $6.17e-01$ |
| Glaucoma vs. PTG | -4.36 | $1.28e-05$ | $7.66e-05$ |
| Healthy vs.PTG (post-diagnosis) | 2.48 | $1.32e-02$ | $7.93e-02$ |
| Glaucoma vs.PTG (post-diagnosis) | -0.26 | $7.99e-01$ | 1.0 |
| PTG vs.PTG (post-diagnosis) | 3.02 | $2.51e-03$ | $1.50e-02$ |

**Supplementary Table 3:** Dunn’s test comparing **PEF** for Glaucoma, healthy, PTG and PTG (post-diagnosis) groups, with Bonferroni correction for p-values.

| Groups | Kruskal-Wallis Z | P (unadjusted) | P (adjusted) |
| --- | --- | --- | --- |
| Healthy vs. Glaucoma | 7.68 | $1.54e-14$ | $9.25e-14$ |
| Healthy vs. PTG | -2.17 | $3.02e-02$ | $1.81e-01$ |
| Glaucoma vs. PTG | -5.12 | $3.01e-07$ | $1.81e-06$ |
| Healthy vs.PTG (post-diagnosis) | 5.58 | $2.38e-08$ | $1.43e-07$ |
| Glaucoma vs.PTG (post-diagnosis) | 2.65 | $8.12e-03$ | $4.86e-02$ |
| PTG vs.PTG (post-diagnosis) | 5.70 | $1.19e-08$ | $7.17e-08$ |

**Supplementary Table 4:** Dunn’s test comparing **Age** for Glaucoma, healthy, PTG and PTG (post-diagnosis) groups, with Bonferroni correction for p-values.

| Groups | Kruskal-Wallis Z | P (unadjusted) | P (adjusted) |
| --- | --- | --- | --- |
| Healthy vs. Glaucoma | −19.26 | $1.10e - 82$ | $6.61e - 82$ |
| Healthy vs. PTG | −5.90 | $3.53e - 09$ | $2.12e - 08$ |
| Glaucoma vs. PTG | 1.47 | $1.41e - 01$ | $8.46e - 01$ |
| Healthy vs. PTG (post-diagnosis) | −11.51 | $1.16e - 30$ | $6.96e - 30$ |
| Glaucoma vs. PTG (post-diagnosis) | −4.15 | $3.33e - 05$ | $2.00e - 04$ |
| PTG vs. PTG (post-diagnosis) | −4.12 | $3.73e - 05$ | $2.24e - 0$ |

**Supplementary Table 5:** Dunn’s test comparing **BMI** for Glaucoma, healthy, PTG and PTG (post-diagnosis) groups groups, with Bonferroni correction for p-values.

| Groups | Kruskal-Wallis Z | P (unadjusted) | P (adjusted) |
| --- | --- | --- | --- |
| Healthy vs. Glaucoma | −6.09 | $1.09e - 09$ | $6.57e - 09$ |
| Healthy vs. PTG | 1.34 | $1.80e - 01$ | 1.0 |
| Glaucoma vs. PTG | 3.69 | $2.26e - 04$ | $1.36e - 03$ |
| Healthy vs. PTG (post-diagnosis) | 1.70 | $8.84e - 02$ | $5.30e - 01$ |
| Glaucoma vs. PTG (post-diagnosis) | 4.05 | $5.07e - 05$ | $3.04e - 04$ |
| PTG vs. PTG (post-diagnosis) | 0.27 | $7.89e - 01$ | 1.0 |

**Supplementary Table 6:** Dunn’s test comparing **BMI** for Glaucoma, healthy, PTG and PTG (post-diagnosis) groups, with Bonferroni correction for p-values.

| Groups | Kruskal-Wallis Z | P (unadjusted) | P (adjusted) |
| --- | --- | --- | --- |
| Healthy vs. Glaucoma | −14.31 | $1.98e - 46$ | $1.19e - 45$ |
| Healthy vs. PTG | −10.74 | $6.32e - 27$ | $3.79e - 26$ |
| Glaucoma vs. PTG | −5.20 | $1.96e - 07$ | $1.18e - 06$ |
| Healthy vs. PTG (post-diagnosis) | −2.40 | $1.64e - 02$ | $9.84e - 02$ |
| Glaucoma vs. PTG (post-diagnosis) | 0.25 | $7.99e - 01$ | 1.0 |
| PTG vs. PTG (post-diagnosis) | 2.50 | $1.23e - 02$ | $7.38e - 02$ |

## 2 Supplementary Methods

**Supplementary Figure 8:**
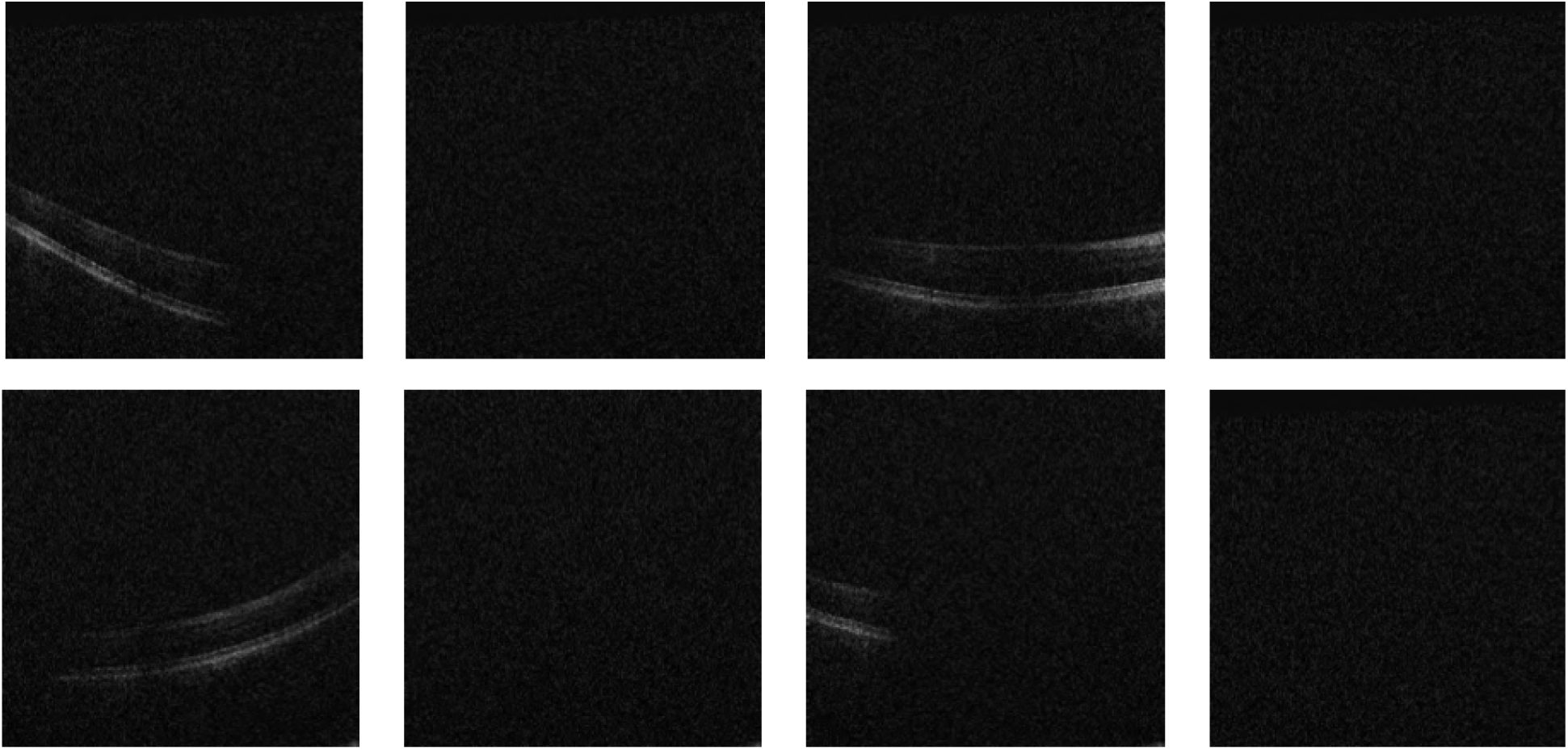
Sample of OCT volumes eliminated due to alignment errors, this shows first two OCT slices for four retinas where alignment failure occured. All OCT slices were eliminated for such retinas.

**Supplementary Figure 9:**
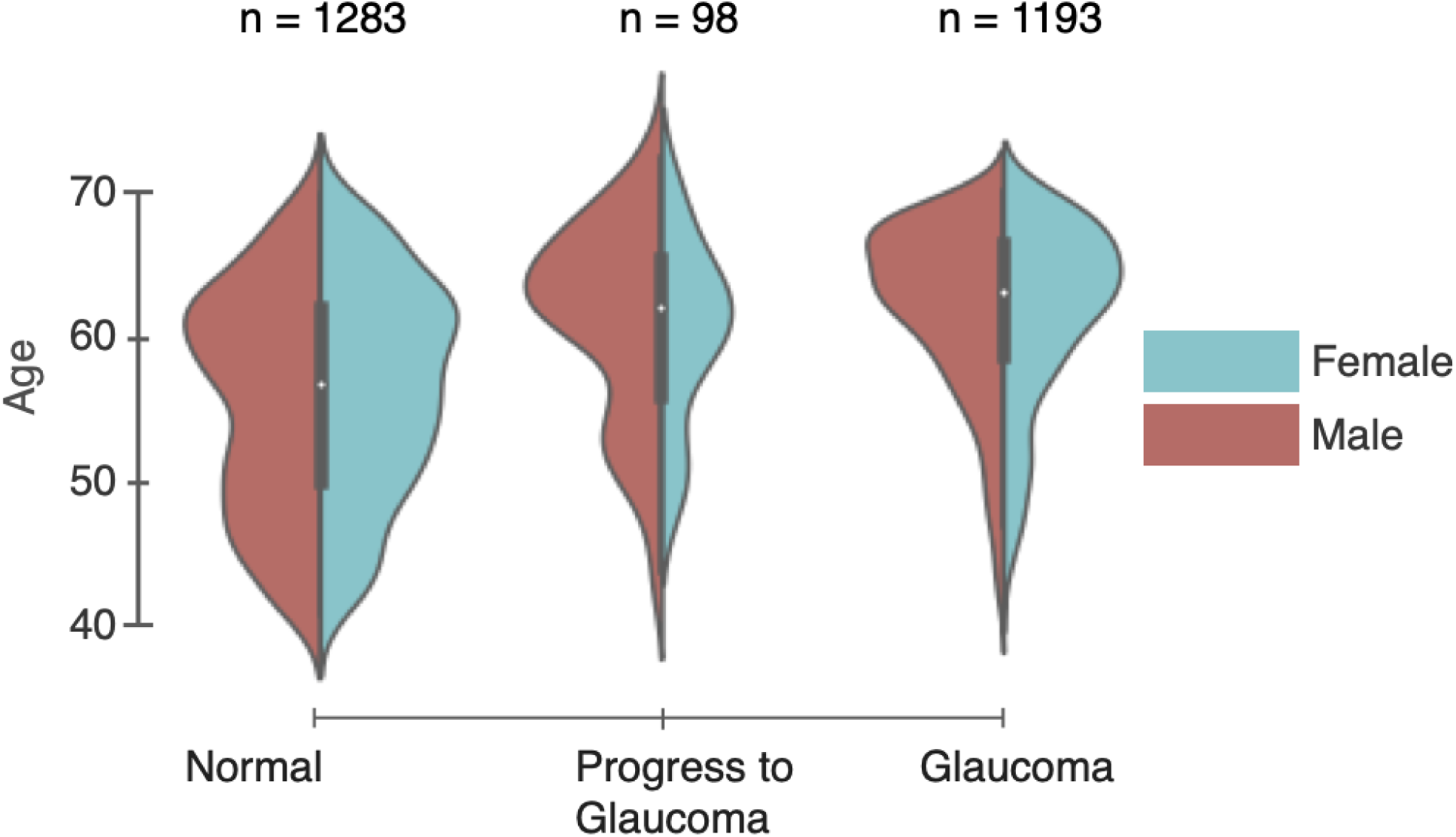
Data distribution: Age and gender distributions as well as number of subjects in the Normal (left), progress to glaucoma (middle) and glaucoma (right) subject groups.

**Supplementary Figure 10:**
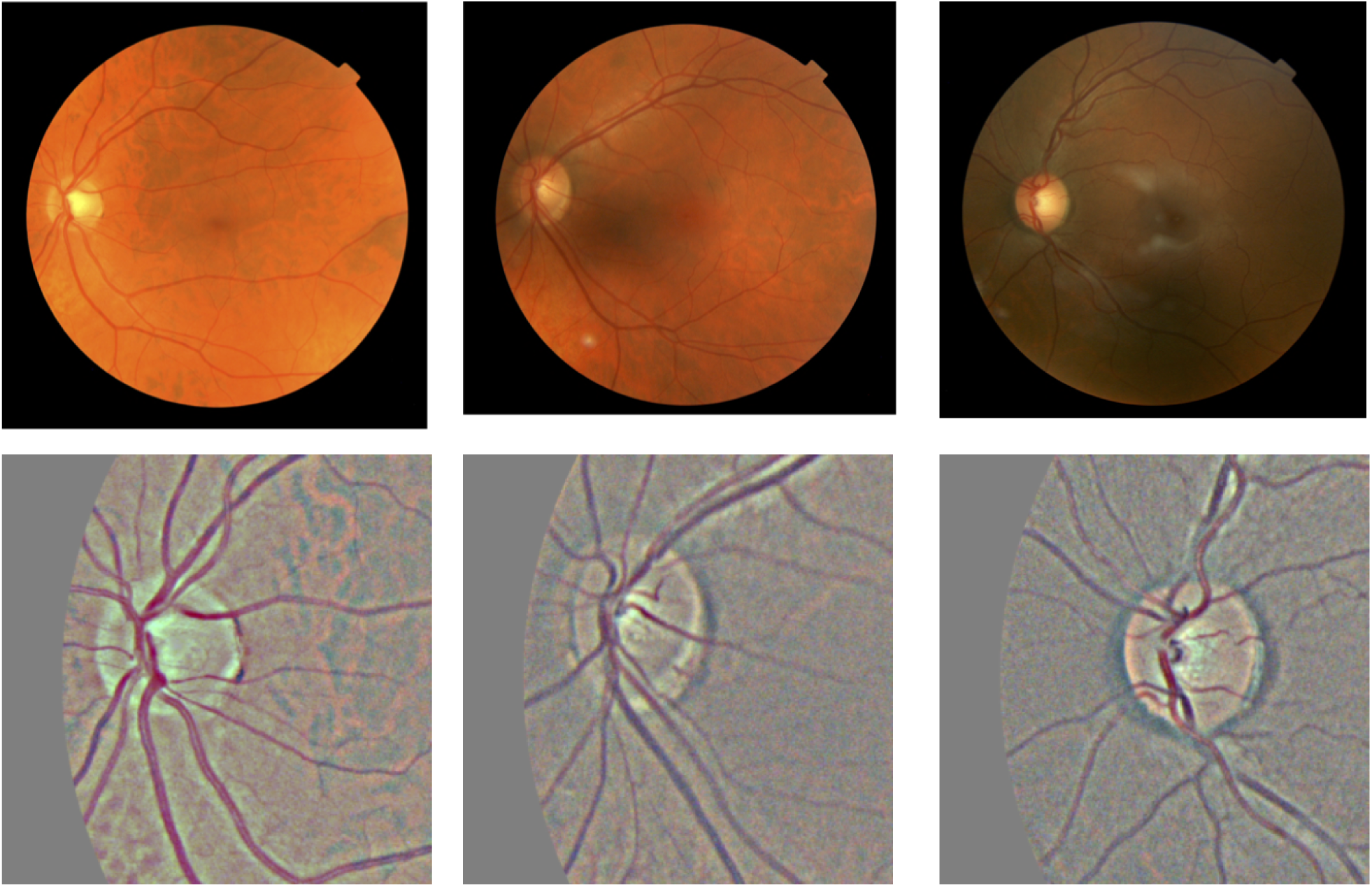
Data processing for CFP images. Top row shows original images, bottom row shows cropped and pre-processed images.

**Supplementary Figure 11:**
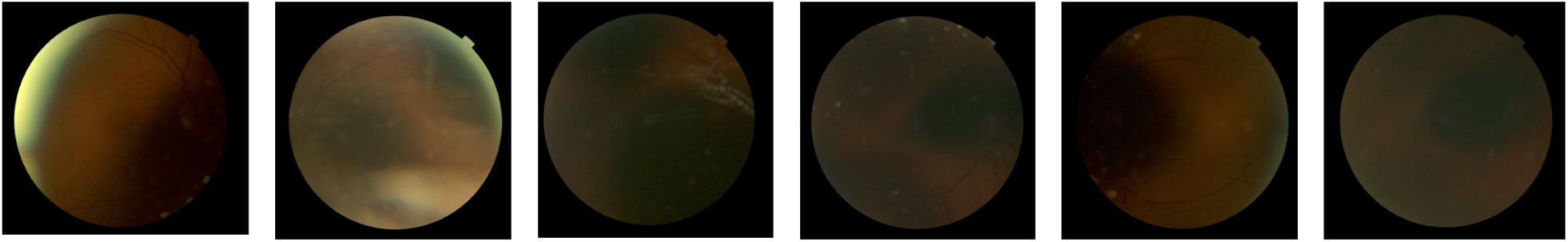
Sample of CFP eliminated from test set due to bad quality.

## Notes

https://github.com/uwdb/automatedglaucomadetection

